# The breathing cycle gates memory reactivation during human NREM sleep

**DOI:** 10.64898/2026.07.29.741474

**Authors:** Esteban Bullón Tarrasó, Tobias Staudigl, Thomas Schreiner

## Abstract

Sleep-dependent memory consolidation relies on the precise coordination of slow oscillations (SOs) and spindles. Respiration has recently emerged as a potential pacemaker of this coordination, but whether it thereby shapes memory processes during sleep remains unknown. Here, we used closed-loop targeted memory reactivation (TMR) in humans (N = 25) to target auditory cues to either each participant’s SO–spindle-favored respiratory phase (preferred-phase cueing) or the opposite phase (antiphase cueing). Preferred-phase cueing enhanced associative memory relative to antiphase cueing and promoted more precise spindle alignment to SO up-states. Respiratory phase also shaped the timing of reactivation: preferred-phase cues aligned category-specific reactivation with the ensuing SO up-state, whereas antiphase cues delayed reactivation to later SO–spindle complexes occurring at a similar respiratory phase. These findings identify breathing as an endogenous timing signal that organizes the SO–spindle windows in which memory reactivation unfolds, thereby structuring sleep-dependent consolidation.

## Introduction

Sleep supports the consolidation of newly acquired memories by promoting the reactivation of recently encoded information during non-rapid eye movement (NREM) sleep^1,2^. Memory reactivation is thought to be organized by a hierarchy of sleep oscillations, in which cortical slow oscillations (SOs), thalamocortical spindles, and hippocampal sharp-wave ripples provide temporally coordinated windows for memory processing^3–5^. SOs delineate alternating periods of widespread neuronal inhibition (down-states) and excitation (up-states) across cortical and subcortical networks^6–8^. Sleep spindles preferentially nest within excitatory SO up-states, forming SO–spindle complexes that create transient windows of enhanced cortical excitability and synaptic plasticity^9–11^. Within these SO–spindle windows, hippocampal ripples are thought to coordinate memory reactivation, thereby supporting the redistribution of memory traces from the hippocampus to neocortical networks^12–14^. Consistent with this framework, more precise spindle alignment to SO up-states has been associated with stronger memory reactivation^15^ and better post-sleep memory retention, linking this temporal coordination to the efficacy of consolidation^16–18^.

These findings point to temporal precision as a key factor in sleep-dependent memory processing. Yet, how the sleeping brain coordinates this oscillatory hierarchy across distributed networks to support memory reactivation remains incompletely understood. Although spindles preferentially cluster around SO up-states^15,19–21^, their timing relative to the SO cycle varies substantially across events, individuals, and development^16,17^. This variability suggests that SO–spindle coupling may be biased by an additional physiological timing signal. In this context, respiration has emerged as a compelling candidate; breathing rhythmically modulates neural excitability and oscillatory activity across widespread brain regions^22–26^, and has been linked to cognitive performance during wakefulness^27–30^. Recent work has extended these observations to human sleep, showing that respiratory phase systematically biases the timing of SOs, spindles, and their interplay in the form of SO–spindle complexes during NREM sleep^31–33^. Critically, however, this evidence remains largely correlational, leaving open whether respiratory phase merely tracks sleep-oscillatory dynamics or actively shapes SO–spindle coupling and memory reactivation, thereby influencing sleep-dependent memory retention.

Here, we used a closed-loop targeted memory reactivation (TMR^34–36)^ paradigm to directly test the functional role of respiratory phase in sleep-dependent memory processing. First, participants learnt associations between auditory cues and images of either objects or scenes. Then, the previously learnt cues were delivered during NREM sleep toward each participant’s individual SO–spindle-favored respiratory phase (i.e., the respiratory phase at which SO–spindle tend to cluster; in-phase cueing) or toward the opposite phase of the respiratory cycle (antiphase cueing). In-phase cueing enhanced sleep-dependent memory retention relative to antiphase cueing and promoted more precise spindle alignment to SO up-states. Moreover, respiratory phase also shaped the temporal profile of memory reactivation: following in-phase cueing, category-specific neural evidence emerged in close temporal alignment with the up-state of the elicited SO–spindle complex, whereas antiphase cueing produced a delayed reactivation profile approximately 2 s later. Notably, this delayed reactivation emerged at a respiratory phase similar to that observed following in-phase cueing and coincided with later SO–spindle complexes, suggesting that memory reactivation is governed by respiration-favored SO–spindle states rather than by cue onset alone. Together, these findings support the idea that respiration acts as an endogenous pacemaker that structures SO–spindle coupling, memory reactivation, and sleep-related memory processing.

## Results

Building on previous evidence that respiration may modulate sleep oscillations during NREM sleep^32,33,37^, we asked whether this coupling is functionally relevant for sleep-dependent memory consolidation. To move beyond correlational observations, we combined respiration-locked closed-loop TMR with overnight EEG recordings in 25 participants (Figure 1a). About one week before the main experimental session, participants completed an adaptation nap (one-hour sleep opportunity; mean NREM sleep = 31.48 ± 2.37 min) to determine the SO–spindle-favored respiratory phase. This participant-specific phase estimate was then used to guide auditory TMR cueing during the subsequent overnight session.

**Figure 1.**
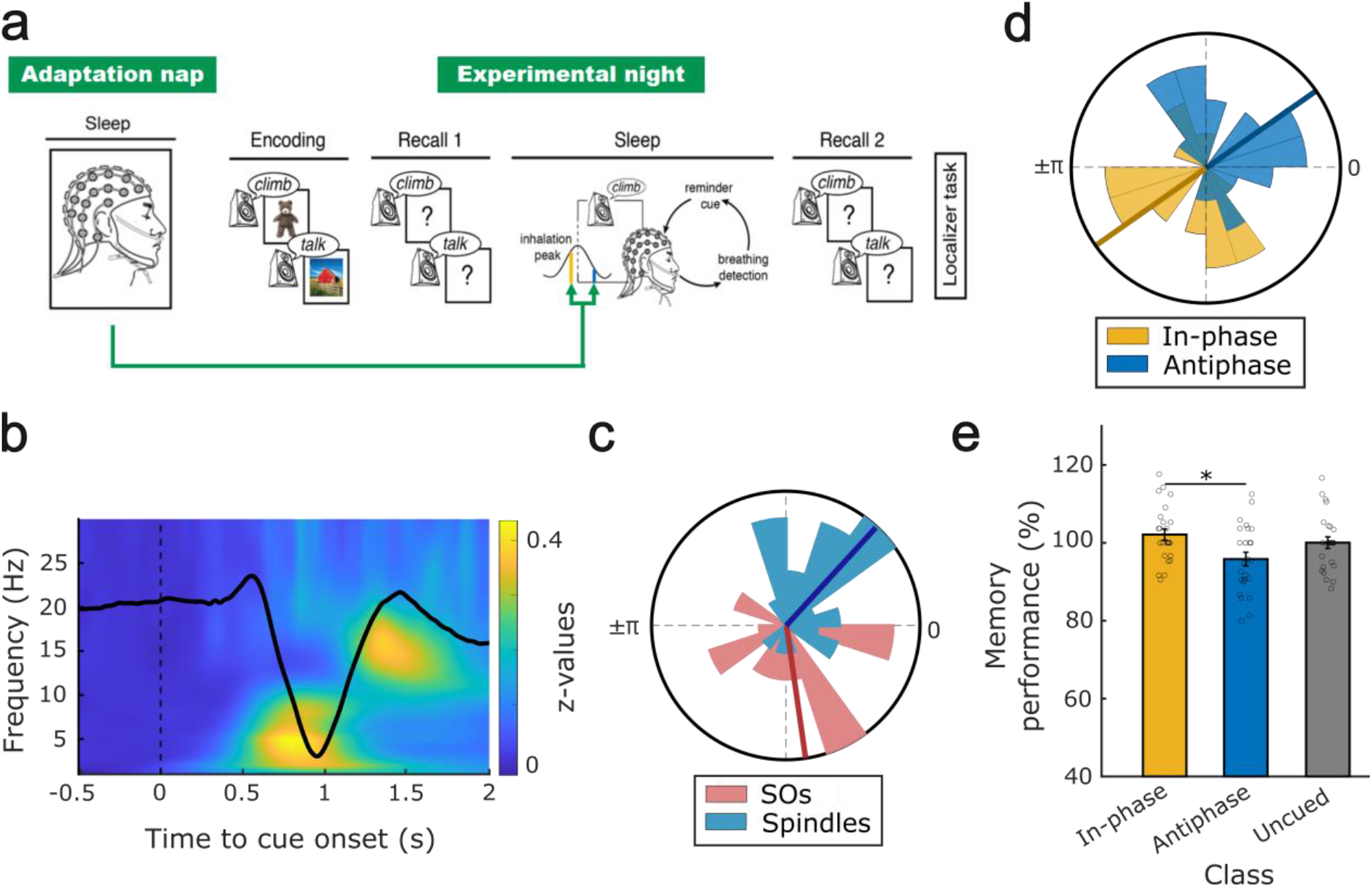
Experimental procedure, TMR-related physiological responses, and behavioral results. **(a)** The experiment began with an adaptation nap, during which each participant’s preferred phase of sleep oscillation coupling to respiration was estimated. One week later, participants completed the main experimental session. During encoding, they learned 90 verb–image associations involving objects and scenes. Memory performance was assessed before and after an overnight period of sleep. During 1 h of NREM sleep, closed-loop TMR was administered by replaying cues at respiratory phases determined from the adaptation nap. At the end of the session, participants completed a localizer task involving a novel set of object and scene images. **(b)** TMR cue-locked EEG responses show a K-complex-shaped event-related potential following cue onset (t = 0), accompanied by increased power in the SO and sigma frequency ranges in the TFR. **(c)** SOs (red) and spindles (blue) are coupled to specific phases of respiration across participants during NREM sleep without TMR. Radial lines represent circular means of each distribution, for electrodes AF8 (SOs, α = −81.978 ± 14.559°) and POz (Spindles, α = 47.584 ± 12.770°). Across participants, SO (z = 6.660) and spindle (z = 8.216) events were non-uniformly distributed across the respiratory cycle (Rayleigh test, both *p* < 0.001). **(d)** Respiratory phases at which cues were delivered for the in-phase (yellow) and antiphase (blue) conditions across participants. Radial lines represent circular means of each distribution (in-phase: α = −144.976 ± 17.059; antiphase: α = 35.025 ± 17.059). Phase distributions were significantly non-uniform (Rayleigh test, z = 5.046, *p* = 0.005). **(e)** Behavioral results revealed a significant main effect of cueing condition on memory performance (repeated-measures ANOVA, F_2,48_ = 4.222, *p* = 0.018). Bar plots depict mean relative memory performance (± SEM), expressed as post-sleep recall normalized to pre-sleep recall, separately for in-phase (yellow), antiphase (blue), and uncued (grey) items. Dots represent individual participant values (*N* = 25). Asterisks indicate significant pairwise differences (*p* < 0.05).

During the main experimental session, participants learned to associate spoken verbs with images depicting either objects or scenes, which were tested before and after sleep using an associative memory task (Figure 1a). During the intervening overnight sleep period, previously learned verbs were re-presented during one hour of stable NREM sleep using respiration-locked closed-loop TMR. Cues were delivered either ∼500 ms before each participant’s preferred respiratory phase (in-phase) or ∼500 ms before the opposing phase (antiphase), while a third set of items remained uncued. This allowed us to target time windows expected to differ in their likelihood of engaging SO–spindle dynamics linked to memory reactivation^15,32^. Cue presentation elicited the expected K-complex-shaped event-related potential (ERP) together with increases in slow-oscillation (<2 Hz) and sigma-band power (12–16 Hz) in the time-frequency representation (TFR)^38,39^ (Figure 1b; see Supplementary Figure 1 for condition-specific ERPs and TFR). After sleep, participants completed an independent object-versus-scene localizer task, which was used to train classifiers for the assessment of category-specific memory reactivation during sleep^15,40,41^.

### Validation of respiration - sleep oscillation coupling

To validate the physiological rationale of the closed-loop cueing protocol, we first asked whether respiration–sleep oscillation coupling during the main experimental night was consistent with prior work^32^. We therefore analyzed 90-min periods of the overnight recording that did not contain TMR (no-TMR periods immediately following TMR), providing an independent estimate of endogenous respiration-linked sleep dynamics (Figure 1c; see Supplementary Figure 2 for TFR and MI replications of ref.^32^). In line with our previous study, respiratory phase modulated low-frequency activity in the SO range (<2 Hz; z > 2.58, *p* < 0.01) and sigma band (12–16 Hz; z > 2.58, *p* < 0.01; Supplementary Figure 2). Time–frequency representations time-locked to inhalation peaks showed significantly greater power in the SO- and sigma-range power than those time-locked to exhalation troughs (*p* = 0.029 and *p* < 0.001, respectively). Event-based analyses further confirmed that SOs, spindles, and SO–spindle complexes were systematically coupled to specific phases of the respiratory cycle: SOs (α = −81.978 ± 14.559°; Figure 1c) and SO–spindle complexes (α = −22.478 ± 15.944°; Supplementary Figure 2) occurred preferentially before inhalation peaks (respective Rayleigh tests: z = 6.660, *p* < 0.001; z = 5.688, *p* = 0.003), whereas spindle events (α = 47.584 ± 12.770°; Figure 1c) occurred thereafter (z = 8.216, *p* < 0.001). We next used these no-TMR periods to validate the participant-specific phase estimates derived from the adaptation nap. Because the no-TMR data provided a larger, and therefore, more stable estimate of endogenous respiration–sleep coupling, nominal cue labels were harmonized relative to this overnight reference for individual participants when necessary (see Methods). This procedure yielded the participant-specific cue assignments shown in Figure 1d, with in-phase cues falling before the inhalation peak in most participants. On average, the circular offset was approximately π/2 relative to the previously reported preferred SO–spindle phase^32^, consistent with the 500 ms phase advance introduced to account for auditory cue delivery and subsequent neural processing.

### Respiratory phase during TMR shapes memory consolidation

Memory performance was assessed before and after sleep as associative memory accuracy (i.e., the proportion of correctly retrieved images), separately for in-phase (102.12 ± 1.45%; mean +/-SEM), antiphase (95.84 ± 1.74%), and uncued (100.05 ± 1.47%) items. To quantify sleep-dependent changes in memory, post-sleep performance was normalized to pre-sleep performance, yielding a relative memory change index (100% indicating no change across sleep). A repeated-measures ANOVA revealed a significant main effect of cueing condition on relative memory change (F_2,48_ = 4.222, *p* = 0.018). Post hoc paired-sample t-tests showed that memory retention was significantly higher for in-phase than for antiphase items (t_24_ = 2,775, *p* = 0.008; Figure 1e), whereas comparisons involving uncued items did not reach significance (in-phase vs. uncued: t_24_ = 1.003, *p* = 0.321; antiphase vs. uncued: t_24_ = −1.850, *p* = 0.070). These findings indicate that the respiratory phase at which TMR cues were delivered modulated the behavioral impact of cueing during sleep (for a full overview of TMR cueing effects on memory performance, see Supplementary Table 3). By contrast, recognition memory showed no significant effect of cueing condition (repeated-measures ANOVA: F_2,48_ = 1.63, *p* = 0.203), suggesting that the phase-dependent behavioral effect was primarily expressed in associative recall rather than recognition memory.

### Respiratory phase during TMR modulates SO–spindle coupling and the timing of memory reactivation

We next asked whether cueing at different phases of the respiratory cycle influenced the occurrence and coordination of sleep oscillations following auditory stimulation during NREM sleep. First, we detected sleep events in the TMR data using established procedures^15,21,36^ (see Methods) and quantified their occurrence relative to cue onset (peri-event time histograms; PETH). Cue-locked analyses of SOs, spindles, and SO–spindle complexes revealed no significant differences in event rates between the in-phase and antiphase conditions (Supplementary Figure 3; all *p* > 0.05), indicating that respiratory phase did not alter the overall likelihood of inducing these oscillations.

We then examined whether respiratory phase might influence the temporal coordination between SOs and spindles, rather than the induction of sleep oscillations per se, given prior evidence that respiration shapes the precise coupling between these oscillations during NREM sleep^32^. To test this, we quantified time-resolved phase–amplitude coupling (PAC; see Methods^42^) between the phase of SOs (0.3–1.5 Hz) and the amplitude of spindle-band activity (12–16 Hz) for each cueing condition. As illustrated in Figure 2a, PAC was significantly stronger in response to in-phase as compared to antiphase cues (*p* = 0.031; 1.273 to 2.070 s from cue onset; corrected across time and electrodes), with the effect most pronounced over frontal electrode sites and centered around SO up-states (see Supplementary Figure 1). To further characterize this effect, we constructed peri-event phase histograms by binning spindle occurrences according to SO phase within the time window and electrode cluster showing significant PAC effects. This analysis revealed a higher proportion of spindles occurring around SO up-states, defined as 0°, in the in-phase condition relative to the antiphase condition (−16.5 to 16.5°, *p* = 0.009, corrected across phase; Figure 2b), indicating that cueing at the preferred respiratory phase promoted the precise alignment of sleep spindles with SO up-states. Moreover, the mean spindle rate of the significant cluster predicted memory performance (β = 2.602, *p* = 0.021; Supplementary Figure 4), regardless of cue class.

**Figure 2.**
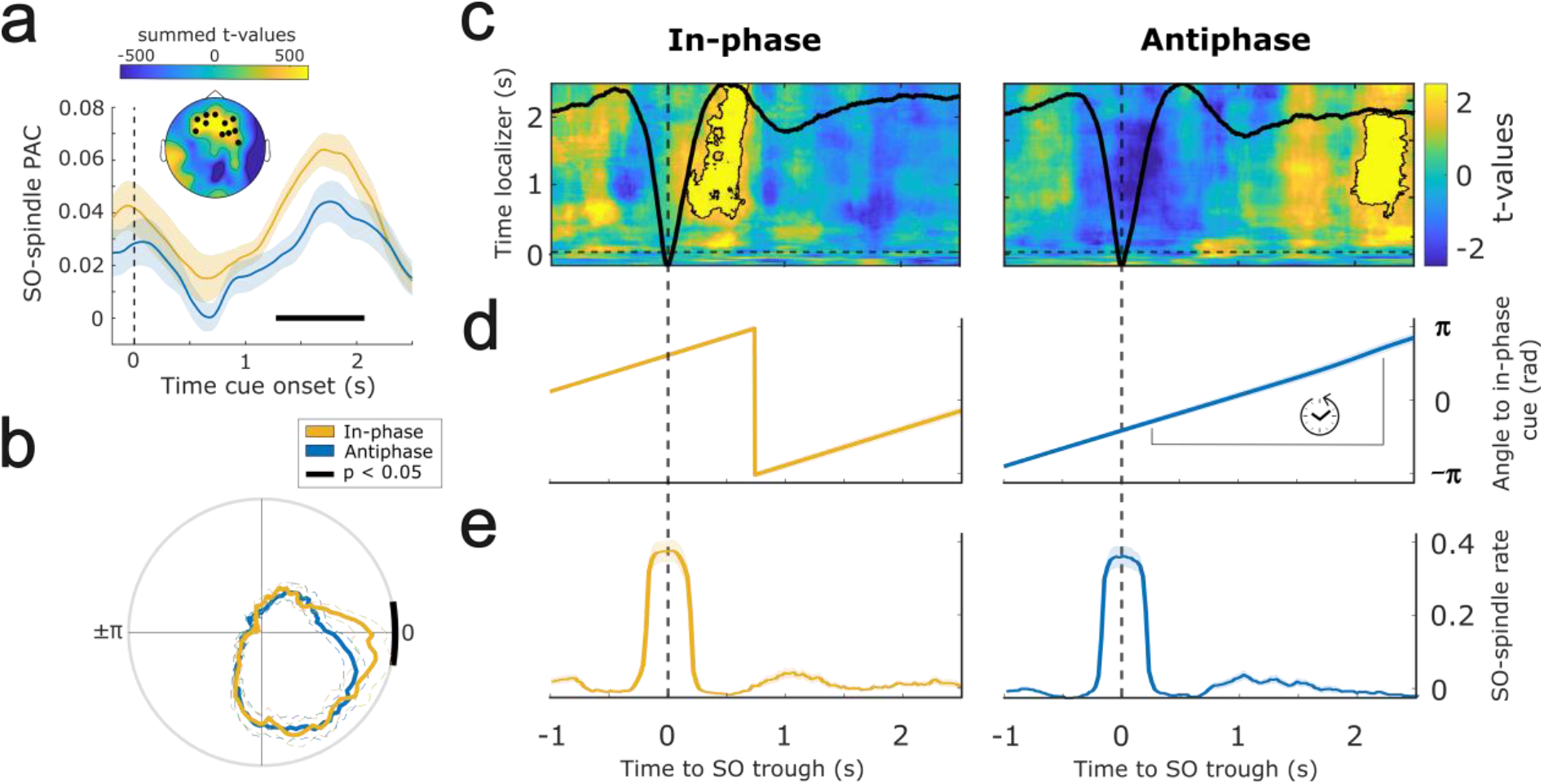
Respiratory phase during TMR modulates SO–spindle coupling and the timing of memory reactivation. **(a)** Phase-Amplitude Coupling (PAC) between SO phase and spindle-band amplitude, time-locked to cue onset, shows increased coupling for in-phase cues (yellow) compared to antiphase cues (blue). The black horizontal bar indicates time intervals with significant differences (*p* = 0.031, 1.273 to 2.070 s from cue onset, corrected for multiple comparisons across time and electrodes). The vertical dashed line marks cue onset (t = 0 s). The inset displays the topography of the significant cluster, primarily located over frontal regions. **(b)** Peri-event phase histogram of spindle occurrences relative to the SO phase reveal a higher proportion of spindles occurring during the SO up-state for in-phase cues (yellow) compared to antiphase cues (blue). The black arc denotes phase bins showing significant differences (−16.5 to 16.5°, p = 0.009, corrected across phase bins). **(c)** Category-specific neural patterns (objects vs. scenes) were significantly decoded above chance from SO–spindle complex–locked EEG data during consolidation for both in-phase (left; 0.160 to 0.691 s, p = 0.042) and antiphase (right; 1.965 to 2.461 s, p = 0.032) conditions. Contour lines indicate significant clusters (corrected across time), and the color scale (blue to yellow) represents *t*-values relative to a null distribution of decoding performance. **(d)** Average respiratory phase difference relative to each participant’s in-phase cue angle and **(e)** PETH of SO–spindle complexes for in-phase (left, yellow) and antiphase (right, blue) conditions.

Having established that respiratory phase modulates SO–spindle coordination, we next asked whether this modulation was accompanied by differences in TMR-triggered memory reactivation during sleep. To this end, we first identified SO–spindle complexes that occurred after TMR cues associated with items successfully recalled before sleep (17.47% of trials, mean of 55.52±6.51 trials per participant). We then extracted EEG epochs from −1 to 2.5 s relative to the SO-trough of each complex. A classifier trained on the independent localizer dataset (−0.2 to 2.5 s relative to stimulus onset; see Supplementary Figure 5) was then applied to these SO–spindle complex–locked EEG segments. This allowed us to test whether EEG activity during TMR contained category-specific information about the cued memories, distinguishing object-from scene-associated cues. Decoding performance was evaluated separately for in-phase and antiphase TMR conditions and compared against condition-specific null distributions generated by label shuffling (object vs. scene categories). In the in-phase condition, decoding performance was significantly above chance within a time window centered on the SO up-state (0.160 to 0.691 s from SO trough), indicating that category-specific memory reactivation emerged in close temporal alignment with the SO–spindle complex (*p* = 0.042, corrected across time, Figure 2c). By contrast, category-specific decoding in the antiphase condition exceeded the corresponding null distribution only in a later time window, approximately 2 s after the initial SO–spindle complex trough (1.965 to 2.461 s from SO trough, *p* = 0.032, corrected across time), rather than around the ensuing up-state (see Supplementary Figure 5b for the direct comparison between conditions). Thus, both cueing conditions were associated with category-specific memory reactivation, but at markedly different times relative to the initially elicited SO–spindle complex.

### Memory reactivation converges on a common respiratory phase and SO–spindle state

Although above-chance classification emerged later in the antiphase condition, the event-related respiratory phase trajectories suggested that significant above-chance classification in both cueing conditions appeared to occur at a similar phase of the respiratory cycle (Figure 2d). This raised the possibility that memory reactivation was more closely tied to respiratory phase than to elapsed time after cue onset. If so, the respiratory phases associated with above-chance decoding should converge across the in-phase and antiphase conditions despite the temporal offset between their reactivation windows. To test this hypothesis, we extracted, for each participant and cueing condition, the mean respiratory phase corresponding to the respective time window of significant above-chance classification (Figure 3a). We then quantified within-participant differences by computing the circular phase difference between in-phase and antiphase reactivation windows. A V-test against 0 rad showed that these phase differences were significantly concentrated around zero (V = 17.701, *p* < 0.001; Figure 3a), consistent with memory reactivation in the two cueing conditions converged on a common phase of the respiratory cycle despite their temporal offset. Together, these findings suggest that memory reactivation is preferentially expressed during respiration-favored phases rather than exclusively at a fixed latency after cue onset.

**Figure 3.**
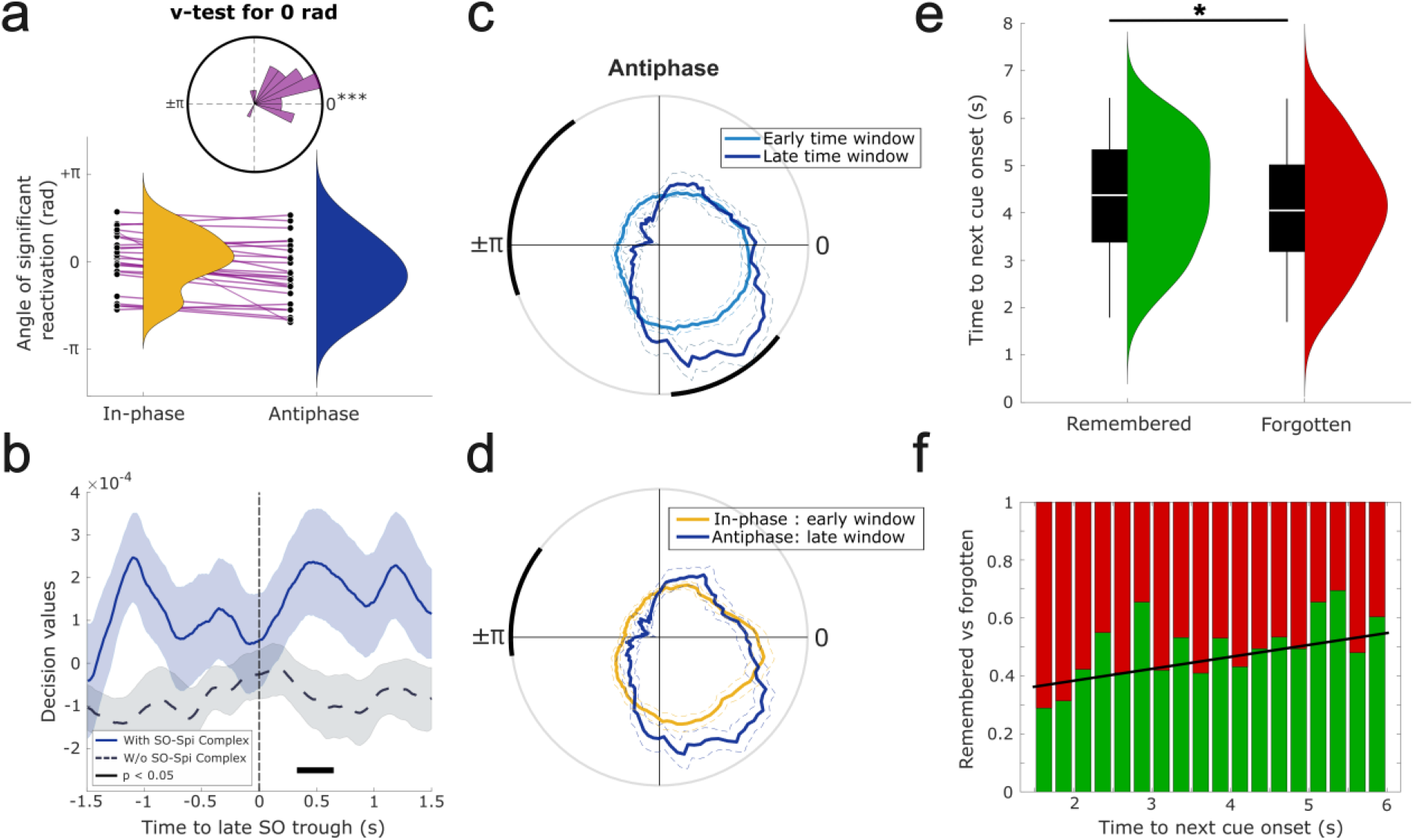
Memory reactivation aligns with a common respiratory phase and SO–spindle dynamics. **(a)** Respiratory phase during periods of significant above-chance classification is similar for in-phase (yellow) and antiphase (blue) cueing within participants. Purple lines connect paired values for each participant. The inset shows a non-uniform distribution of phase differences centered around 0 rad, as confirmed by a *V*-test (V = 17.701, *p* < 0.001). **(b)** Category-specific neural patterns (objects vs. scenes) were successfully decoded from EEG data time-locked to SO–spindle complexes occurring in a later post-cue window (1.5–3.5 s), compared to trials without such complexes in this interval (0.328 to 0.648 s from SO trough, *p* = 0.040, corrected across time). **(c)** SO–spindle complexes occurring at a later time window (dark blue) present a stronger spindle coupling than the elicited ones in the original time window of interest (light blue) during antiphase trials. Black arc lines represent phases of significant difference (down-state cluster: p = 0.001; up-state cluster: p = 0.012). **(d)** Late window SO–spindle complexes (same as in c) present a similar coupling of spindles in the up-state of SOs as compared to SO–spindle complexes elicited by in-phase cues (yellow). **(e)** For antiphase trials, correctly recalled items were associated with longer intervals between the late SO–spindle complex and the onset of the subsequent cue compared to forgotten items, as determined by a generalized linear mixed-effects model (β = 0.230, p = 0.041). **(f)** Probability of successful recall as a function of the interval between the late SO–spindle complex and the next cue onset in the antiphase cueing condition. Green (red) bars indicate the proportion of remembered (forgotten) items within each time bin. The black line illustrates the corresponding increase in recall (and decrease in forgetting) probability across time (for visualization purposes).

The convergence in respiratory phase raised the question of whether delayed antiphase reactivation was also embedded in the same oscillatory state as preferred-phase reactivation. We therefore tested its alignment with later SO–spindle complexes. To test this directly, we re-epoched antiphase trials around SO–spindle complexes occurring after the initial post-cue response window (23.99±1.61% out of all previously remembered trials; 1.5–3.5 s after cue onset). The previously trained classifier was then applied to these later SO–spindle complex–locked segments, yielding a single-trial, time-resolved metric that reflects the classifier’s confidence in assigning a specific category^40,43^ (i.e., decision values; see Methods). The resulting decision values were compared with those from trials lacking such late SO–spindle complexes. Decoding performance was significantly above chance in this analysis (0.328 to 0.648 s from SO trough, *p* = 0.040, corrected across time; Figure 3b), indicating that delayed antiphase reactivation was again expressed around SO up-states when trials were aligned to later, endogenous SO–spindle complexes.

We then tested whether, within the antiphase condition, later SO–spindle complexes showed stronger SO–spindle coordination than those occurring within the initial post-cue time window. As shown in Figure 3c, spindles were more tightly coupled towards SO up-states in later than earlier antiphase complexes (*p* = 0.012; −85.5 to −37.5°; corrected across phase; see Supplementary Figure 6a for a similar comparison within in-phase cues). Next, we assessed whether these late antiphase complexes resembled the SO–spindle state observed after in-phase cueing. To test directly whether the preferred coupling phases were aligned across conditions, we calculated the within-participant circular difference between the mean coupling angles for in-phase and late-antiphase complexes and applied a V-test with an expected direction of 0 rad. The phase differences were significantly concentrated around zero (V = 15.275, *p* < 0.001; Supplementary Figure 6b), indicating that SO–spindle coupling in the two conditions exhibited similar preferred phase relationships.

Finally, we asked why antiphase cueing, despite being followed by detectable signatures of memory reactivation when later SO–spindle complexes occurred (similar to the in-phase memory reactivation profile), nevertheless resulted in poorer memory performance than in-phase cueing. Prior work has shown that auditory stimulation delivered shortly after a TMR cue can abolish the memory benefits of TMR, consistent with the idea that reactivation-related plasticity requires a protected post-cue window^44,45^. We therefore reasoned that the delayed reactivation observed in the antiphase condition may have placed reactivation-related processing closer to the onset of subsequent cues, given the minimum inter-cue interval of 5 s (see Methods), thereby increasing its susceptibility to interference. To test this, we fitted a generalized linear mixed-effects model (GLMM) predicting post-sleep recall performance (modeled as a binomial outcome) as a function of the interval between the late SO–spindle complex and the onset of the subsequent cue. Participant and item were included as random effects (see Methods). The analysis revealed a significant positive effect of the interval duration, such that longer delays between reactivation and the next cue were associated with higher recall probability (β = 0.230, *p* = 0.041; Figure 3e). To further illustrate this relationship, we estimated forgetting probability as a function of the interval between the late SO–spindle complex trough and the onset of the subsequent cue. As shown in Figure 3f, successful retrieval probability increased monotonically with increasing interval duration (*r* = 0.619, *p* = 0.006; Pearson’s correlation), suggesting that delayed reactivation on antiphase trials remained vulnerable to disruption when the subsequent cue occurred too soon (see also ref.^45^).

## Discussion

The current study identifies the breathing cycle as an endogenous timing signal that gates memory reactivation during human NREM sleep. Using closed-loop TMR, we show that cues delivered toward each participant’s SO–spindle-favored respiratory phase enhanced associative memory retention compared with cueing near the opposing respiratory phase. This behavioral benefit was accompanied by more precise spindle alignment to SO up-states and a phase-dependent shift in the timing of memory reactivation: whereas preferred-phase cueing aligned category-specific reactivation with the ensuing SO up-state, antiphase cueing delayed reactivation until a later SO–spindle complex occurred at a similar respiratory phase. Together, these findings suggest that respiration structures sleep-dependent memory consolidation by shaping the SO–spindle windows in which memory reactivation can effectively unfold.

These findings provide experimental support for an emerging framework in which respiration coordinates offline memory processing across species^33^. In rodents, breathing has been shown to organize cortico–hippocampal dynamics, including hippocampal ripples^46^. Extending these findings to human NREM sleep, we previously showed that respiration modulates SOs, spindles, and SO–spindle coupling, and that the strength of respiration–SO–spindle coupling is correlated with the extent of memory reactivation during SO–spindle complexes^32^. Here, we build directly on this work by moving from an observational link between respiration and sleep-dependent reactivation to a direct test of whether respiratory phase shapes the SO–spindle windows in which memory reactivation unfolds and supports post-sleep memory. By cueing toward versus away from each participant’s SO–spindle-favored respiratory phase, we show that respiratory timing modulates SO–spindle precision, the temporal dynamics of category-specific memory reactivation, and post-sleep memory performance. Mechanistically, respiratory phase did not appear to determine whether auditory cues elicited SOs, spindles, or SO–spindle complexes, as cue-locked event rates were comparable across conditions (Supplementary Figure 3). This likely reflects the strong capacity of auditory cues to evoke sleep-oscillatory responses^47^ even outside the respiration-favored phase. Instead, respiratory phase shaped the timing of cue-evoked SO–spindle responses, such that preferred-phase cueing promoted more precise spindle alignment to SO up-states (Figure 2a, b). This temporal precision has been shown to be functionally important because spindles nested in SO up-states coincide with increased cortical excitability and plasticity-promoting conditions, creating physiological windows for memory reactivation and strengthening^10,11,15,48^. Consistent with this interpretation, the respiratory phase also shaped when memory reactivation emerged. Following preferred-phase cueing, category-specific reactivation was closely aligned with the SO up-state of the associated SO–spindle complex (Figure 2c, left). Thus, preferred-phase cueing coordinated memory reactivation with the same oscillatory state in which spindles were most precisely aligned to the SO up-state.

The antiphase condition provides a critical extension and test of this interpretation. Antiphase cueing did not simply abolish reactivation. Instead, robust category-specific reactivation emerged approximately two seconds later (Figure 2c, right), at a respiratory phase similar to preferred-phase reactivation (Figure 2d, Figure 3a). This delayed reactivation was again linked to SO–spindle dynamics: when antiphase trials were realigned to later SO–spindle complexes, category-specific reactivation re-emerged around SO up-states (Figure 3b). This suggests that the late reactivation for the antiphase condition was not simply a delayed cue-locked response. Instead, it occurred within a later SO–spindle state, in which spindles were aligned to the SO up-state to a similar degree as after preferred-phase cueing (Figure 3c, d). Together, this pattern suggests that respiration shapes when robust memory reactivation emerges, by biasing the timing of SO–spindle states that support its expression. Alternatively, antiphase cues might have initially triggered weaker or less detectable memory-related activity, with category-specific reactivation becoming measurable using scalp EEG only once a later, more favorable SO–spindle state emerged. In both cases, memory reactivation during sleep appears not to be purely locked to external cue onset, but to depend on endogenous respiratory and SO–spindle timing. Future intracranial recordings with single-unit activity could test whether antiphase cues initially engage memory-specific neuronal ensembles only weakly, or whether such ensemble reinstatement emerges only later within a more favorable respiratory–SO–spindle state.

In either case, the state-dependent delay may carry a functional cost that helps explain the behavioral deficit observed in the antiphase condition. We found that delayed reactivation in antiphase trials occurred closer to the onset of the subsequent TMR cue, and that this interval predicted post-sleep recall (Figure 3e, f). This finding suggests that effective TMR does not depend on a single reactivation event. Instead, cue-triggered memory processing may unfold across successive endogenous consolidation windows. This interpretation is consistent with rodent studies showing that auditory cues can bias subsequent replay and that cue-related activity can persist for several seconds to influence later memory-related activity^49,50^ as well as with human work indicating repeated reactivation following a single TMR cue^51^. In addition, prior work in humans suggests that reactivation-induced plasticity requires an undisturbed post-cue window of at least 1–1.5 seconds to be effective; when this window is disrupted by incoming information, the mnemonic benefit is abolished^44,45^. In the antiphase condition, delayed reactivation may therefore leave too little time for such recurrent processing before the next cue occurs. Thus, antiphase cueing may remain capable of eliciting detectable reactivation, but this reactivation may be less likely to translate into a behavioral benefit because it is temporally compressed and more vulnerable to interference by subsequent stimulation.

Overall, these findings may have broader implications for the hierarchical dialogue between the hippocampus and neocortex. While our recordings were limited to scalp EEG, the present results raise the possibility that respiratory timing also structures hippocampal components of sleep-dependent memory reactivation. Hippocampal ripples are thought to coordinate information transfer between the hippocampus and neocortex during sleep^14,21,52,53^, and prior work suggests that respiration can organize interactions among SOs, spindles, and ripples in rodents and humans^37,46,54^. An important future direction will be to examine this tripartite coupling directly in humans using intracranial EEG to test whether respiratory phase coordinates hippocampo–cortical memory reactivation within the SO–spindle–ripple hierarchy.

Together, our findings identify respiration as a physiological timing signal that shapes the SO–spindle windows in which memory reactivation unfolds during human NREM sleep. By linking respiratory phase to SO–spindle precision, category-specific reactivation, and post-sleep memory retention, the present study extends models of sleep-dependent consolidation beyond a purely brain-intrinsic account of oscillatory coordination. Breathing thus emerges as an additional bodily rhythm that helps coordinate the neural events underlying memory processing during sleep. This perspective may inform future closed-loop approaches that incorporate respiratory phase to more precisely target favorable consolidation windows. It may also have clinical relevance, as conditions characterized by disrupted breathing, such as obstructive sleep apnea, could impair memory in part by disturbing the temporal coordination of sleep oscillations and memory reactivation.

## Methods

### Participants

A total of 36 volunteers were initially recruited for the study. Of these, two participants withdrew following the adaptation nap session, and one additional participant was not invited to the main experimental session due to insufficient sleep data obtained during the nap. Furthermore, five participants were excluded due to issues arising during data collection, including technical errors (n = 3), medical reasons (n = 1), and predominant mouth breathing during sleep (n = 1). Following data inspection, an additional three participants were excluded: one due to poor overall data quality and two due to the absence of a discernible cue-evoked event-related potential (ERP) in the EEG. The final sample thus comprised 25 participants (mean age: 24.00 ± 3.37 years; 17 females) included in the analyses. All participants were German native speakers, reported no history of respiratory or neurological disorders, including sleep apnea, and provided written informed consent prior to the experimental procedures. Pre-study screening included several questionnaires: the Pittsburgh Sleep Quality Index (PSQI)^55^, the Morningness-Eveningness Questionnaire^56^, and a custom questionnaire assessing general health and stimulant use. None of the participants reported taking medication at the time of testing, and all were free of neurological and psychiatric disorders. All participants indicated generally good sleep quality and had not engaged in night-shift work for at least eight weeks prior to the experiment. Participants were instructed to abstain from alcohol the evening before the session and from caffeine on the day of testing. Risk for sleep-disordered breathing was assessed using the STOP-BANG questionnaire (Snoring, Tiredness, Observed apneas, high blood Pressure, BMI > 35 kg/m², Age > 50, Neck circumference > 40 cm, male Gender)^57^, which yields scores ranging from 0 to 8; scores ≥ 3 indicate an increased risk of obstructive sleep apnea. All participants in the present sample scored below 3, reflecting a low risk of sleep-disordered breathing. In combination with the absence of increased awakenings in sleep scoring and the lack of observed breathing cessations during visual inspection of respiratory data, these findings suggest that undiagnosed sleep-disordered breathing was unlikely to have affected the results. Participants received either course credit or financial compensation for their participation. The study protocol was approved by the ethics committee of the Department of Psychology at Ludwig-Maximilians-Universität München.

### Procedure

#### Habituation nap

One week prior to the main experimental night, participants completed an adaptation nap in the sleep laboratory. This session served two primary purposes: first, to habituate participants to the laboratory environment and thereby minimize potential first-night effects during the main experimental session; and second, to determine each participant’s preferred phase of sleep oscillation–respiration coupling. This individualized phase estimate was subsequently implemented during the main experimental night within the closed-loop TMR paradigm. The nap session was scheduled at a time of day chosen by the participant, with the recommendation to select a period during which they anticipated increased sleepiness. To facilitate sleep onset, participants were instructed to curtail their habitual sleep duration by approximately one hour on the preceding night and to abstain from caffeine and alcohol on the day of the nap. The nap opportunity lasted 60 minutes (mean NREM sleep = 31.480 ± 2.372 min), during which respiration and 16-channel EEG (F3, F1, Fz, F2, F4, FC1, FCz, FC2, C3, Cz, C4, O1, Oz, O2, M1, M2) were continuously recorded.

#### Preferred phase computation

Following the adaptation nap, each participant’s preferred respiratory phase was determined using the following procedure. First, the respiratory phase corresponding to each detected SO and SO–spindle complex (defined as an SO followed by a spindle within 1.5 s of the SO trough) was extracted from periods of N2 and N3 sleep for each frontal and fronto-central electrode. For each event type and electrode, circular statistics were computed, and phase non-uniformity was assessed using the Rayleigh test, yielding both a z-statistic and associated *p*-value. The preferred phase was defined as the circular mean of the SO phases in the electrode exhibiting a significant phase distribution with the highest z-statistic. If no electrode showed a significant phase preference for SOs, the same procedure was applied to SO–spindle complexes. If neither event type yielded a significant phase distribution, the preferred phase was determined as the circular mean of the phases from the electrode and event type associated with the highest z-statistic overall. To derive the cueing phase for the targeted memory reactivation (TMR) protocol during the main experimental night, the participant-specific preferred phase was adjusted to account for stimulus delivery and neural processing delays. Specifically, the mean respiratory frequency during N2 and N3 sleep was estimated for each participant, and a phase advance corresponding to 500 ms was added to the preferred phase. This adjusted phase was then used as the target phase for closed-loop auditory cue presentation during sleep.

#### Experimental task

The main experimental session commenced at approximately 20:00 with the preparation and placement of EEG electrodes and the respiration sensor. During the encoding task, participants learned 90 verb–image associations, in which auditorily presented German verbs were paired with images of objects or scenes displayed on a computer screen. Memory performance was assessed both before and after sleep. In addition, a functional localizer task was administered at the end of each session (see Localizer Task below). Stimulus presentation and behavioral response collection were controlled using Psychophysics Toolbox Version 3 implemented in MATLAB R2018b (MathWorks, Natick, MA, USA).

#### Stimuli

The stimulus set comprised 300 German verbs and 240 images, evenly divided into object and scene categories, presented across the experimental sessions. Object images depicted animals, food items, clothing, tools, or household objects against a white background (e.g., a calculator), whereas scene images represented identifiable landscapes or locations (e.g., an island). These categories were selected because they reliably engage distinct neural systems, including the Lateral Occipital Complex for object processing^58^ and the Parahippocampal Place Area for scene processing^59^. This categorical dissociation enabled the investigation of experience-dependent memory reactivation during retrieval (see Multivariate Analysis below). All images were drawn from the database published by Konkle et al. (2010).

#### Familiarization

The experiment began with an image familiarization phase designed to facilitate subsequent verb–image associative learning and to establish standardized image labels for later cued recall. Each trial commenced with a fixation cross (1.5 ± 0.1 s), followed by the presentation of one of 90 images that were later used in the encoding phase and were accompanied by captions that accurately labeled the depicted exemplar.

#### Encoding

Each encoding trial began with a fixation cross (1.5 ± 0.1 s), followed by the auditory presentation of a German verb via loudspeakers (e.g., “essen”). The corresponding image was then displayed for 4 s. Participants were instructed to form a vivid mental image or narrative linking the verb to the depicted object or scene. After image offset, participants indicated whether the constructed mental representation was realistic or bizarre. Participants were informed in advance that their memory for the verb–image associations would be tested. The encoding block was repeated twice, with randomized trial order in each iteration, to achieve stable memory performance as determined in a prior pilot study.

#### Memory Tests

Memory for the verb–image associations was assessed before and after the sleep interval. Each test included all previously encoded verbs, intermixed with 50 novel verbs (foils). Trials began with a fixation cross (1.5 ± 0.1 s), followed by auditory presentation of a verb. After a 3 s delay, participants indicated whether the verb was “old” (previously learned) or “new” (foil). For “new” responses, the next trial commenced immediately. For “old” responses, participants were prompted to type a description of the associated image or to indicate “do not know” if recall failed. Responses were scored as correct if the typed description either matched the original familiarization caption or unambiguously corresponded to the target image content.

#### Localizer Task

During the localizer task, participants viewed a novel set of 180 images (90 objects, 90 scenes), irrespective of experimental condition. Each trial began with a fixation cross (1.5 ± 0.1 s), followed by presentation of a randomly selected image for a variable duration (2.5 ± 0.1 s). Each image was presented twice. Participants indicated whether the image was being shown for the first time (“new”) or the second time (“old”).

#### Closed-loop Targeted-Memory Reactivation

The sleep period commenced at approximately 23:30, following completion of the pre-sleep memory assessment. Participants were allotted 7–8 h of sleep opportunity (sleep architecture parameters are reported in Supplementary Table 1). During the first hour of stable N2 and N3 sleep, a closed-loop TMR protocol was administered. Previously learned German verbs were replayed auditorily at specific phases of the respiratory cycle, individually determined from each participant’s adaptation nap.

The learned verbs were divided into three experimental conditions: (1) in-phase cues, presented at the participant’s preferred phase of sleep oscillation–respiration coupling; (2) antiphase cues, delivered at the opposite respiratory phase (i.e., preferred phase ± π radians); and (3) uncued items, which served as a no-stimulation control condition. In addition, 30 task-irrelevant control verbs were presented during sleep (15 in-phase, 15 antiphase) to control for nonspecific auditory stimulation effects. To ensure balanced baseline performance across conditions, item assignment was adjusted based on pre-sleep memory accuracy, resulting in an equivalent distribution of correctly recalled items across the three experimental conditions. In-phase and antiphase cues were delivered in pseudorandomized order during stable NREM sleep. The TMR procedure was implemented using OpenViBE Brain Computer Interface Software^61^. Respiratory signals were preprocessed online and transformed into phase estimates in real time (see Respiration Preprocessing below). Auditory cues were triggered when the ongoing respiratory signal reached the predefined target phase and at least 5 s had elapsed since the preceding cue. Throughout stimulation, the experimenter continuously monitored polysomnographic signals to adjust cue volume (40 to 65 dB, with starting loudness of 50 dB) or suspend stimulation if signs of arousal emerged. After a cumulative total of 60 min of TMR had been delivered, cueing was terminated, and participants were allowed to sleep undisturbed until morning awakening.

#### EEG and respiratory recordings

Electrophysiological data were acquired using the eego 65-channel EEG system (ANT Neuro, Enschede, Netherlands). Electrode impedances were maintained below 20 kΩ. EEG signals were referenced online to the CPz electrode and digitized at a sampling rate of 512 Hz. Respiratory activity was recorded using a Braedon cTherm Cannula Thermistor (Braebon Medical Corporation, NJ, USA). The thermistor cannula was positioned beneath the nostrils to detect temperature fluctuations associated with airflow during inhalation and exhalation. To minimize displacement during sleep, the cannula was gently secured to the participant’s cheeks with medical tape. The respiration signal was recorded continuously alongside the EEG using the same acquisition system, ensuring precise temporal synchronization between respiratory and neural recordings.

### Data analysis

#### Behavioral analysis

Memory performance was quantified based on associative recall accuracy. Specifically, performance was calculated as the proportion of correctly recalled images, computed separately for the pre-sleep and post-sleep memory tests. To assess sleep-related changes in memory, post-sleep performance was normalized by pre-sleep performance, yielding a relative memory change index. Values above 100% indicated an improvement in associative memory after sleep, whereas values below 100% indicated a decline. This measure was calculated separately for each of the three TMR conditions: in-phase, antiphase, and uncued items. Differences in sleep-related memory change across the three cueing conditions were evaluated using a repeated-measures analysis of variance (rm-ANOVA). When the significant main effect was observed, post hoc pairwise comparisons were conducted using paired-sample t-tests to determine which conditions differed from each other.

#### EEG and respiration pre-processing

All data were analyzed using MATLAB (2020a; MathWorks). EEG data were preprocessed using the FieldTrip toolbox for EEG/MEG analysis (v.16/07/2023). All data were downsampled to 256 Hz, filtered around 50 Hz to remove line noise (4^th^ order, two-pass Butterworth filter), and re-referenced to the average of the mastoid channels. Noisy EEG channels were identified by visual inspection, discarded, and then interpolated using a weighted average of the neighboring channels. EEG epochs related to the TMR session were extracted between −2 and 5 seconds locked to cue onset. Sleep staging was carried out offline according to standard criteria^62^ by two independent raters.

For analyses related to the respiratory phase, the phase was extracted by first filtering the respiratory trace around ± 0.1 Hz of the peak frequency (two-pass Butterworth filter) and then computing the angle from the Hilbert transform. Thus, an angle of 0° represents the inhalation peak (i.e., the inhalation-to-exhalation transition), whereas an angle of 180° (± π rad in plots) represents the exhalation trough (i.e., the exhalation-to-inhalation transition). In the case of the closed-loop TMR, the respiratory phase was computed in real time. First, the respiratory signal was one-pass filtered between 0.05 and 0.5 Hz (2^nd^ order Butterworth filter). Then, to get the immediate respiratory phase for each subsequent time-point, we extracted the last 16 seconds of the respiratory signal and computed the inhalation peak-to-exhalation trough (or exhalation trough-to-inhalation peak, depending on the current respiratory phase) median duration of the last respiratory cycles (within the mentioned time window). Lastly, by using the current time to the last inhalation peak (or exhalation trough), we interpolated the immediate phase.

#### Event detection

Sleep events (SOs and spindles) were identified for each participant, based on established detection algorithms^15,21,36^. SOs were detected from the preprocessed EEG signal following band-pass filtering between 0.3 and 1.5 Hz. Positive-to-negative and negative-to-positive zero-crossings were then identified. An event was classified as an SO when the interval between two consecutive positive-to-negative transitions ranged between 0.8 and 2 s and the corresponding trough-to-peak amplitude exceeded twice the standard deviation of the signal within the extracted epoch. A cue was considered to be associated with an SO when the SO trough occurred between 0.5 and 1.5 s following cue onset (as the interval of increased rate illustrated in the peri-event time histogram of SO rates, see Supplementary Figure 3) and was detected in at least half of the channels identified as most prominent for SO activity during a separate detection procedure performed on sleep data recorded in the absence of targeted memory reactivation (see Supplementary Figure 2).

Sleep spindles were detected after band-pass filtering the EEG signal between 12 and 16 Hz. The root-mean-square (RMS) envelope of the filtered signal was computed using a sliding window of 200 ms. Spindle events were defined as periods during which the RMS envelope exceeded a threshold corresponding to 50% above the mean envelope amplitude of the analyzed epoch, with a duration between 0.5 and 3 s. A cue was considered to be associated with a spindle if spindle onset occurred at least 0.5 s after cue presentation and the spindle’s maximum amplitude fell within 3 s following cue onset (as the interval of increased rate shown in the peri-event time histogram of spindle rates, see Supplementary Figure 3). This criterion had to be met in at least one of the channels identified as most prominent for spindle activity during detection performed on sleep data without TMR (see Supplementary Figure 2). SO–spindle complexes were detected as the SOs that were followed by a spindle within 1.5 seconds from the SO trough. The number and proportion of cues with an associated sleep event are recorded in Supplementary Table 2.

#### Preferred phase correction

To verify the assignment of TMR cues to the in-phase and antiphase conditions, we re-estimated each participant’s preferred respiratory phase using an independent portion of the overnight recording that was free of TMR stimulation. Specifically, SO–spindle complexes were detected during a 90-min NREM sleep period following the completion of the TMR session, using the same detection procedure described above. The no-TMR data provided a more precise estimate of endogenous respiration–sleep coupling due to the larger quantity of data. The preferred respiratory phase was then computed from these events and compared with the respiratory phases at which in-phase and antiphase cues had been delivered. If the preferred phase estimated from the overnight recording was closer to the phase originally assigned to the antiphase condition than to that assigned to the in-phase condition, the cue labels for that participant were inverted (i.e., all in-phase cues were relabeled as antiphase, and vice versa). According to this criterion, cue labels were swapped for a total of eight participants.

#### Phase-amplitude coupling

Phase–amplitude coupling (PAC) between the phase of SOs and the amplitude of spindle-band activity was quantified (Figure 2a). The EEG signal was first band-pass filtered in the SO range (0.3–1.5 Hz) and in the spindle range (12–16 Hz). The filtered signals were then subjected to a Hilbert transform to extract the instantaneous phase of the SO component and the amplitude envelope of the spindle-band signal. PAC was computed using the estimator proposed by Özkurt & Schnitzler (2011). The analysis was performed in a time-resolved manner using sliding windows of 1 s with a step size of 50 ms. To account for baseline fluctuations, PAC values were normalized by subtracting the mean coupling strength calculated during the 2 s interval preceding cue onset. PAC estimates were calculated separately for each cueing condition (in-phase and antiphase) and for each EEG channel.

#### Peri-event histograms

To assess differences in sleep events elicited by cueing condition (in-phase vs. antiphase), peri-event time histograms (PETHs) were constructed. Target events included SO troughs, spindle time points of minimum amplitude, and SO–spindle complex troughs, expressed relative to cue onset (Supplementary Figure 3) or to SO troughs (Figure 2e). Histograms were computed with a bin size of 50 ms over a time window ranging from −0.2 to 2.5 s for cue-locked PETH (Supplementary Figure 3) and −1 to 2.5 s for SO trough-locked PETH (Figure 2e). To account for differences in overall event rates, histograms were normalized by dividing the number of detected events per bin by the total number of events within the time window of interest. The resulting time courses were smoothed using a 250 ms moving average window and subsequently z-scored relative to a 500 ms pre-cue baseline period. These steps were performed separately for each participant and electrode.

In addition, peri-event phase histograms (Figure 2b, Figure 3c, d) were constructed to characterize the distribution of spindle occurrences across the SO cycle for both in-phase and antiphase conditions. Analysis was restricted to the electrodes and time interval showing significant effects in the PAC analysis. The instantaneous phase of the SO signal (0.3–1.5 Hz) was derived using the Hilbert transform, with the SO upstate defined as 0°. The phase corresponding to each spindle minimum value was extracted and binned using a resolution of π/60 radians. Spindle occurrence rates within each phase bin were normalized relative to the mean spindle rate during the 2 s preceding cue onset. Finally, phase histograms were smoothed using a moving average window spanning nine bins (window size = 3π/20).

#### Multivariate analysis

Multivariate classification analyses of single-trial EEG data were conducted using MVPA-Light. Classification was implemented with Linear Discriminant Analysis (LDA), which has been widely used for time-resolved decoding of electrophysiological signals^15,40,43,63^. For classification of EEG data from the localizer task, signals were first z-scored across trials independently at each time point. The data were then subjected to principal component analysis (PCA) to transform the signal into a set of orthogonal components ranked according to the variance explained by each component. PCA served to reduce dimensionality and limit overfitting by retaining only the first components accounting for 95% of the total variance. To determine whether object and scene representations could be discriminated in the localizer data, a classifier was trained to distinguish between trials containing object images and those containing scene images. Prior to classification, the data were smoothed using a 200 ms running-average window. EEG channels were treated as features, and a separate classifier was trained and tested at each time point. Classification performance was quantified using the area under the receiver operating characteristic curve (AUC), which reflects the probability that a randomly selected pair of trials from the two classes is correctly assigned to their respective categories (0.5 = chance level; 1.0 = perfect classification). To reduce overfitting, a fivefold cross-validation procedure was applied. The cross-validation procedure was repeated five times with different random fold assignments, and the resulting decoding accuracies were averaged across iterations. Statistical significance was assessed using a permutation approach. Specifically, surrogate decoding accuracies were generated through 250 iterations of label shuffling, yielding a null distribution under the assumption of label exchangeability. Results from within-localizer classification can be seen in Supplementary Figure 5a.

To investigate category-specific memory reactivation following TMR cues (Figure 2c, Supplementary Figure 5b), we applied the temporal generalization method. First, cues that were associated with an SO–spindle complex were identified. EEG epochs were extracted from −1 to 2.5 s relative to the trough of the complex elicited by cues, as prior work has demonstrated that memory reactivation during sleep is temporally linked to this event^15^. A classifier was trained at each time point using the localizer data and subsequently applied to each time point within the sleep reactivation epochs. This approach enabled the assessment of whether neural patterns characteristic of object versus scene processing during wakefulness re-emerged during sleep. Decoding analyses were restricted to cues corresponding to previously remembered associations to ensure the presence of a memory trace available for reactivation. Because the training (localizer) and testing (sleep) datasets were independent, cross-validation was not required for this step. Classification performance was again quantified using the AUC metric. For Figure 3b, decision values^43^ were computed for each trial before their alignment to the later SO–spindle complex and averaged across significant Localizer timepoints (Supplementary Figure 5a).

#### Generalized Linear Mixed-Effects Model

To examine whether the temporal interval between memory reactivation and the onset of the subsequent cue influenced post-sleep memory performance, we fitted a generalized linear mixed-effects model (GLMM). Again, we only included cues belonging to remembered trials during the pre-sleep recall task. The dependent variable was post-sleep recall accuracy for each verb, coded as a binary outcome (1 = correctly recalled, 0 = not recalled) and modelled using a binomial distribution with a ‘logit’ link function. The primary predictor was the time interval between the detected late SO–spindle complex and the onset of the next auditory cue. To account for variability across participants and stimuli, random intercepts were included for both participant and verb identity. The model therefore assessed whether longer or shorter intervals between the SO–spindle complex and subsequent cue presentation were associated with differences in recall probability while controlling for item- and participant-level variability. The model can be expressed as:

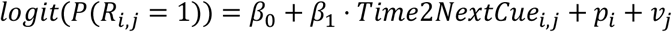

where *R_i,j_* represents the recall outcome for verb *j* in participant *i*, *β_0_* denotes the fixed intercept, *β_1_* represents the fixed effect of the interval between the late SO–spindle trough and the next cue onset, and *p_i_* and *v_j_* correspond to random intercepts for participant *i* and verb *j*, respectively, with *p_i_ ~ N(0, σ^2^_i_)* and *v_j_ ~ N(0, σ^2^_j_)*.

#### Statistical analyses

Behavioral retrieval performance was normalized by computing the ratio of post-sleep to pre-sleep memory accuracy separately for each cueing condition (in-phase, antiphase, and uncued). A repeated-measures analysis of variance (ANOVA) was conducted to assess the effect of cueing condition on relative memory performance. Significant effects were followed up with post hoc paired-sample t-tests comparing all pairs of conditions, which revealed a primary difference between in-phase and antiphase cueing. To test the relation between spindle rate during SO up-states and memory performance, an Analysis of Covariance (ANCOVA) was performed with the spindle rate as co-variate.

Circular data were analyzed using appropriate directional statistics using the Circular Statistics Toolbox^64^. Non-uniformity of phase distributions was assessed using the Rayleigh test without assuming a preferred direction (e.g., Figure 1c, d; Supplementary Figure 2b). In contrast, for analyses testing whether two distributions had a similar mean (reactivation phase for in-phase and antiphase cueing, Figure 3a; spindle rate during SO up-state, Supplementary Figure 6b), a circular mean of 0 rad was specified a priori, and a V-test was applied to the difference between both distributions.

Unless otherwise stated, statistical analyses of time and phase-based data were performed using non-parametric cluster-based permutation tests as implemented in FieldTrip^65^. These tests do not assume a specific underlying data distribution and control for multiple comparisons across time (or phase) points. At the sample level, dependent-samples t-tests were computed to identify clusters of contiguous significant samples across participants, using a threshold of *p* = 0.05. Cluster-level statistics were calculated using the maximum cluster sum (‘maxsum’) of t-values, and statistical significance was determined via Monte Carlo permutation testing (α = 0.05, two-tailed).

## Acknowledgments

We thank Dr. Hong-Viet V. Ngo-Dehning for helpful discussions during the initial development of the closed-loop stimulation protocol. T. Sc. is supported by the Emmy Noether program of the German Research Foundation (492835154).

## Data availability

All data supporting the findings of this study will be made publicly available upon publication.

## Code availability

All code related to the analyses of the manuscript will be made publicly available upon publication.

## Author contributions

E.B.T and T.Sc. conceived the study and designed the experiment. E.B.T conducted the experiment. E.B.T and T.Sc. analyzed the data. E.B.T., T.St. and T.Sc. wrote the paper.

## Supplementary material

**Supplementary Figure 1.**
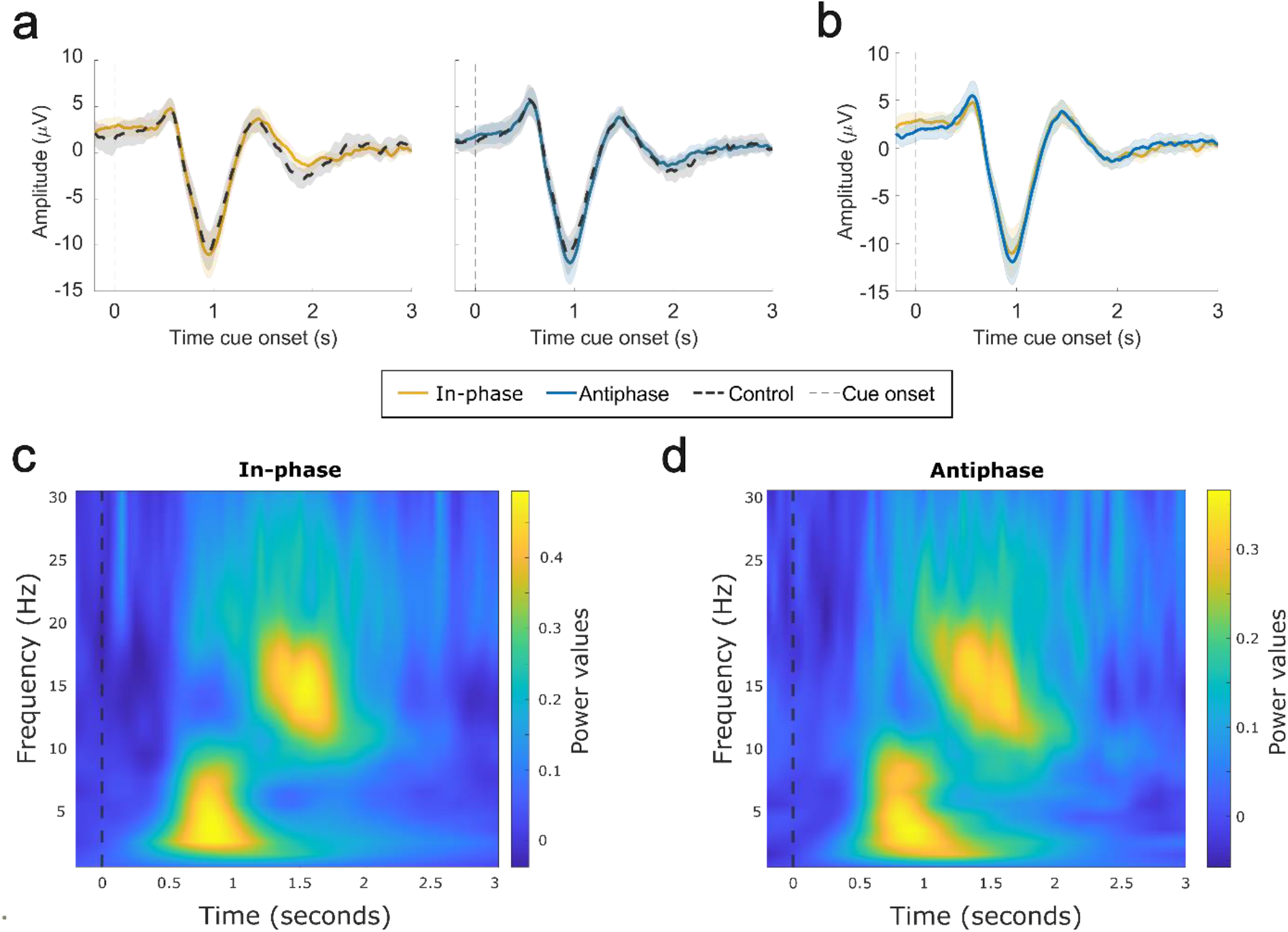
Event-related potentials (ERPs) and time-frequency representations (TFRs) per cue class. **(a)** ERP elicited by memory encoded do not present significant differences in contrast to control cues played in the same phase (left: in-phase; right: antiphase). **(b)** Direct comparison between in-phase and antiphase ERP. **(c, d)** TFR elicited by in-phase (c) and antiphase (d) cues. Cluster permutation test shows no significance between both conditions.

**Supplementary Figure 2.**
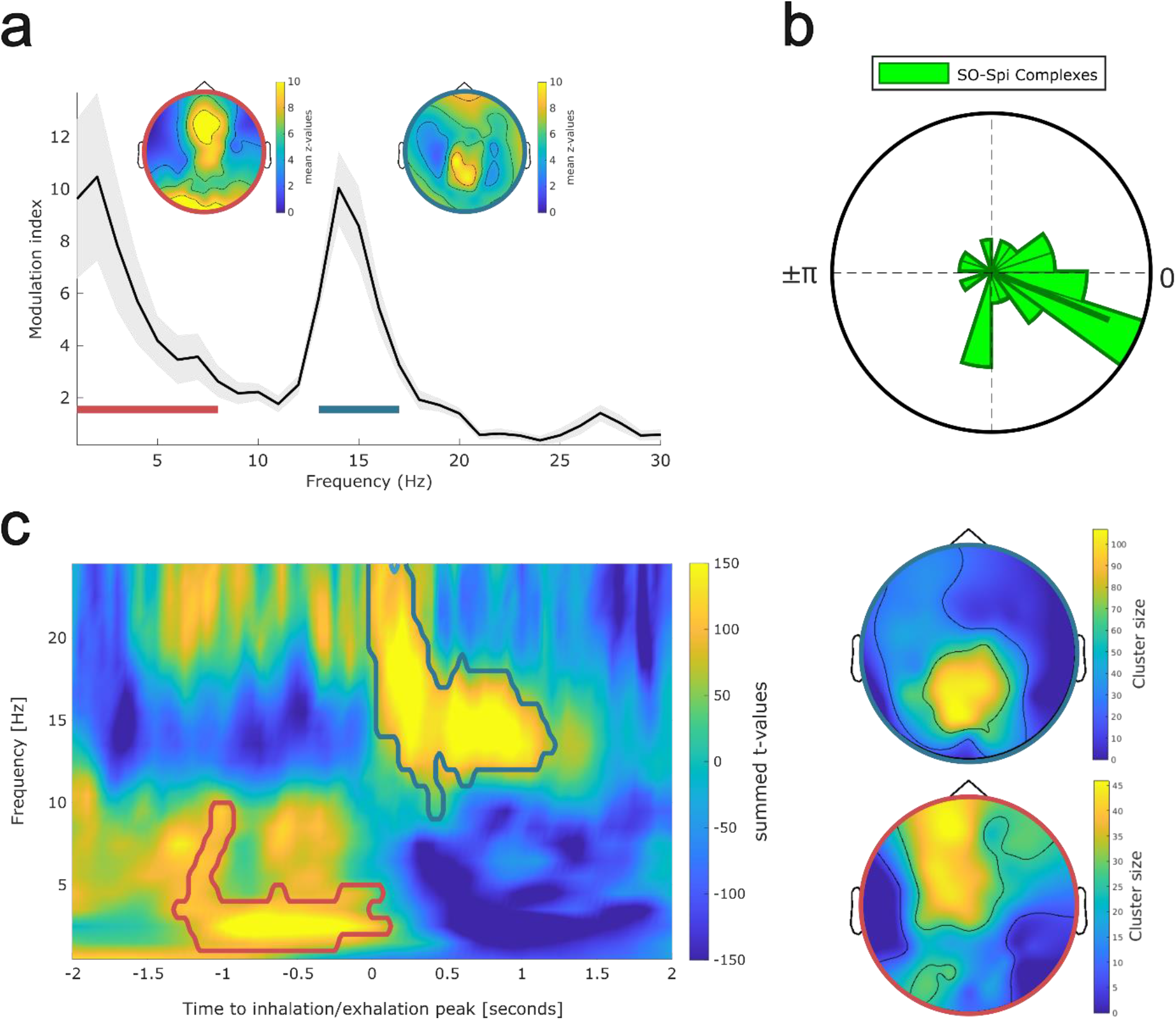
Replication of results presented in Schreiner et al., 2023. **(a)** Modulation index between EEG amplitude and respiratory phase indicates significant modulation in delta (<4 Hz) and sigma (13-17 Hz) centered in the frontal and parietal regions, respectively (z > 2.58, *p* < 0.01, for both bands). **(b)** SO–spindle complexes were non-uniformly distributed across participants in relation to the respiratory phase (Rayleigh test; z = 5.688, *p* = 0.003; electrode FPz). **(c)** Time–frequency representation of NREM sleep EEG data, contrast between inhalation peak and exhalation trough locked segments (mean z-values across significant electrodes). Contours indicate significant clusters in the SO (red, *p* = 0.029) and spindle (blue, *p* < 0.001) ranges (two-sided; corrected across time, frequency and electrodes). The topographies illustrate statistical results across electrodes for each cluster.

**Supplementary Figure 3.**
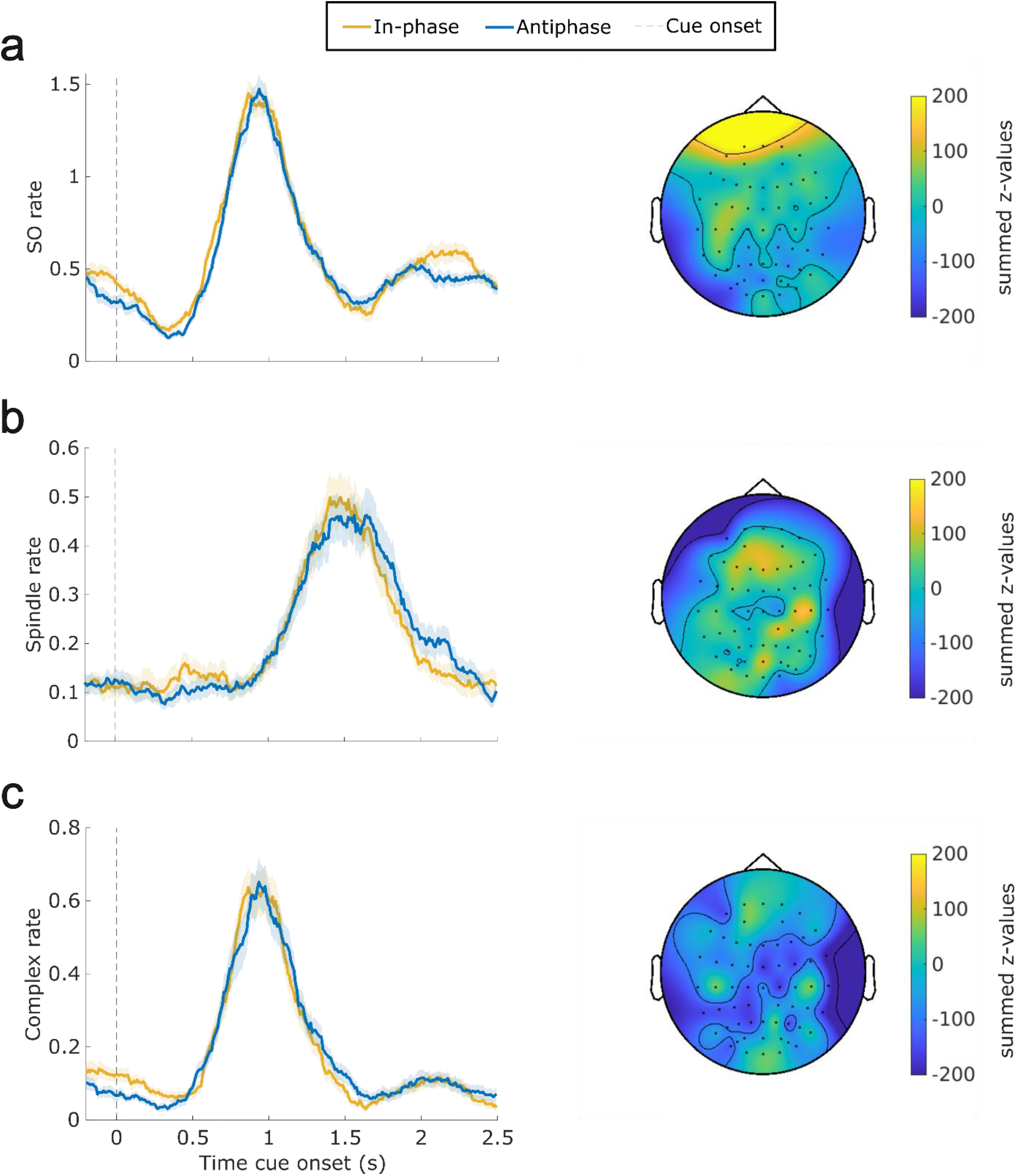
Peri-event time histograms for SOs (a), spindles (b), and SO–spindle complexes (c). There was no significant difference between cue classes (in-phase vs antiphase) for any of the sleep events (two-sided, corrected across time and electrodes). Topographies illustrate the summed z-values across time.

**Supplementary Figure 4.**
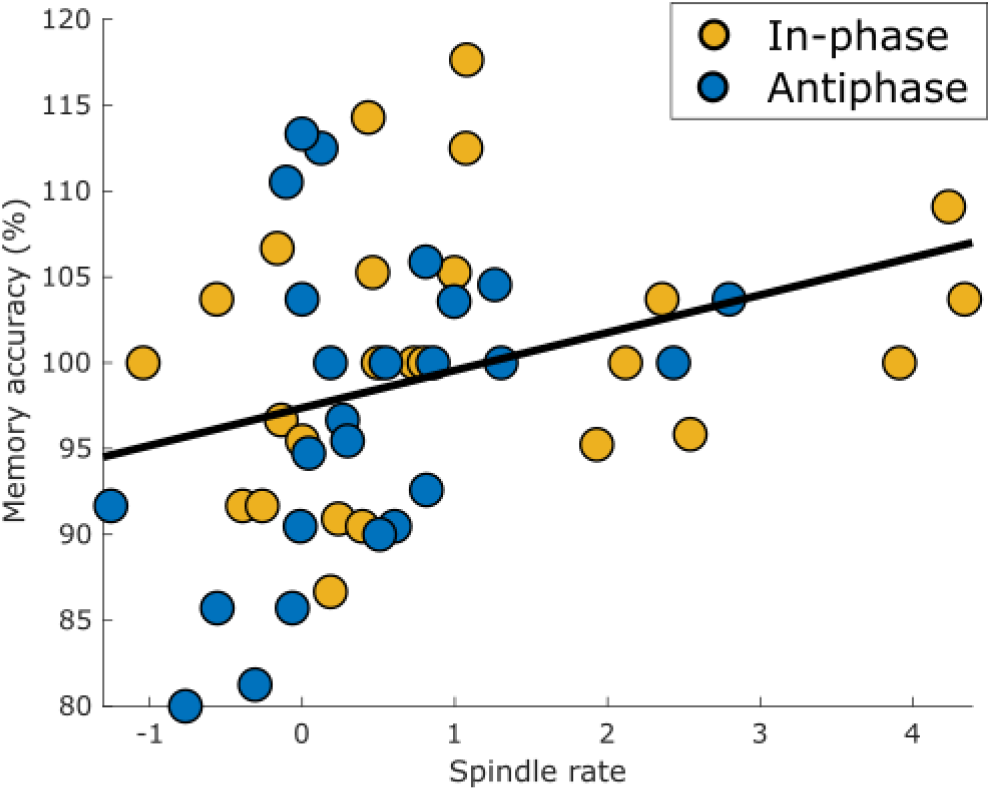
Analysis of covariance between spindle rate and memory accuracy. Spindle rate (specifically during the up-state reaching significance from figure 2a) successfully predicted memory accuracy (β = 2.602, *p* = 0.021; regardless of cue class).

**Supplementary Figure 5.**
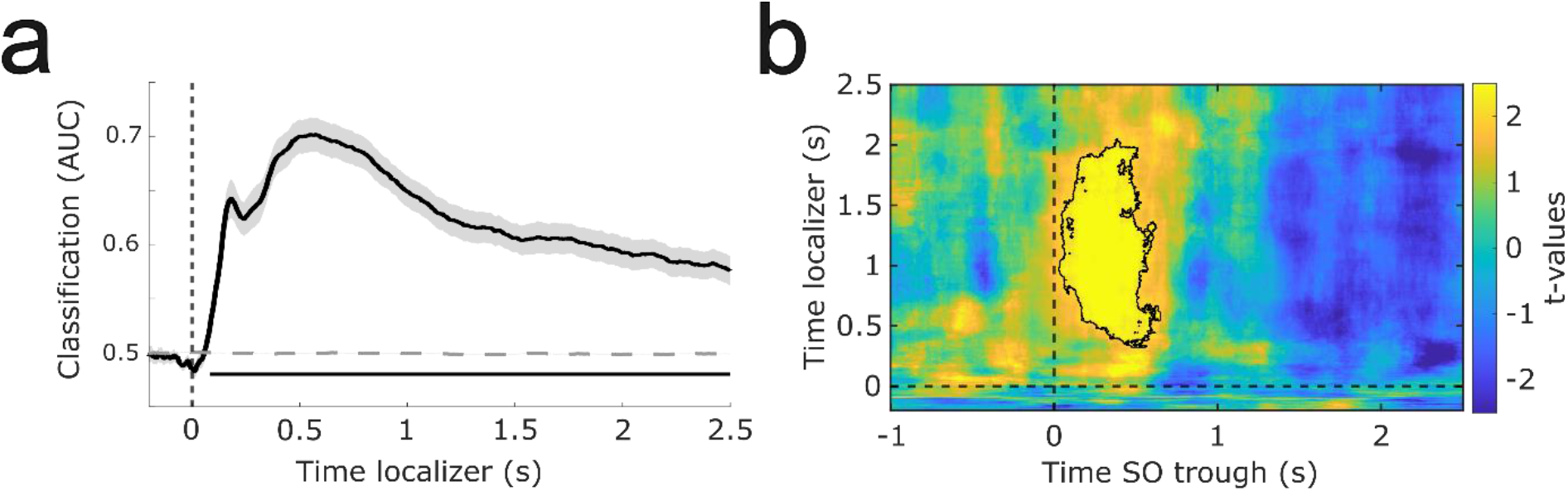
Within localizer classification and in-phase vs. antiphase decodability. **(a)** Stimulus categories (objects vs. scenes) could be decoded above chance from the localizer EEG data, starting around 100 ms post stimulus onset. The black solid line indicates decoding performance (±SEM). The horizontal dashed line indicates surrogate decoding performance, which was estimated by shuffling the training labels 250 times. The vertical dashed line indicates stimulus onset (time = 0 s). The lower horizontal dark grey line shows the temporal extent of significant decoding results as derived from a dependent-samples t-test (two-sided, *p* < 0.001, corrected across time). **(b)** Category-specific neural patterns (objects vs. scenes) were successfully decoded from SO–spindle complex–locked EEG data during consolidation after in-phase compared to antiphase cues (*p* = 0.029, two-sided, corrected across times).

**Supplementary Figure 6.**
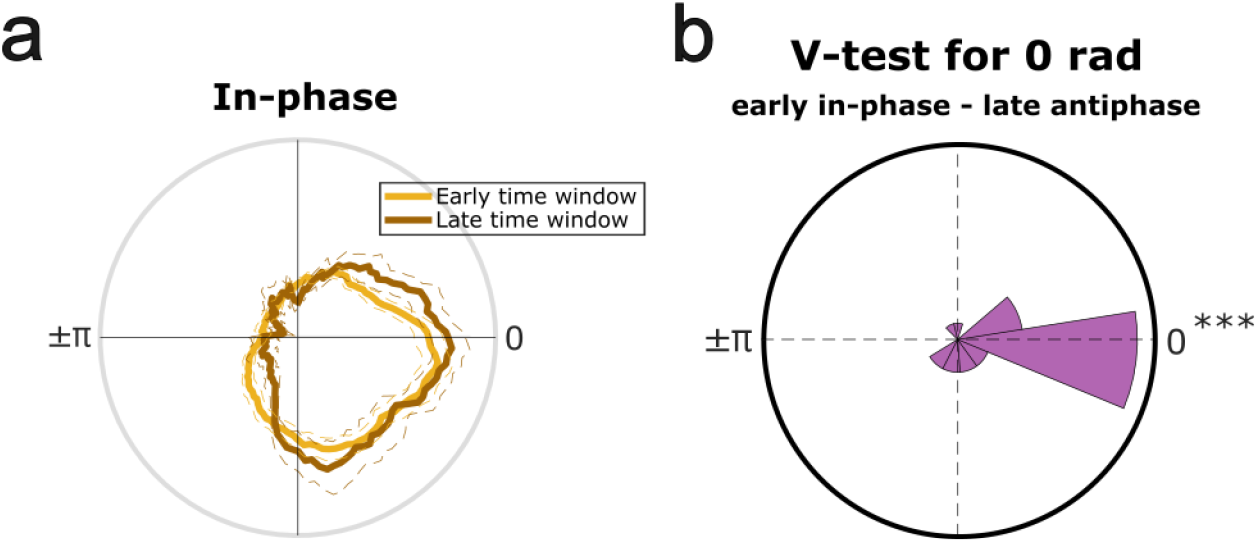
Comparison in SO-spindle coupling between time windows and conditions. **(a)** There is no difference in SO–spindle complexes occurring at a later time window (brown) as compared to the elicited ones in the original time window of interest (yellow; two-sided, corrected across phase). **(b)** Comparison of mean SO–spindle coupling between late antiphase complexes and original in-phase complexes (Figure 3d) revealed no significant differences (V = 15.275, *p* < 0.001).

---

**Supplementary Table 1.**
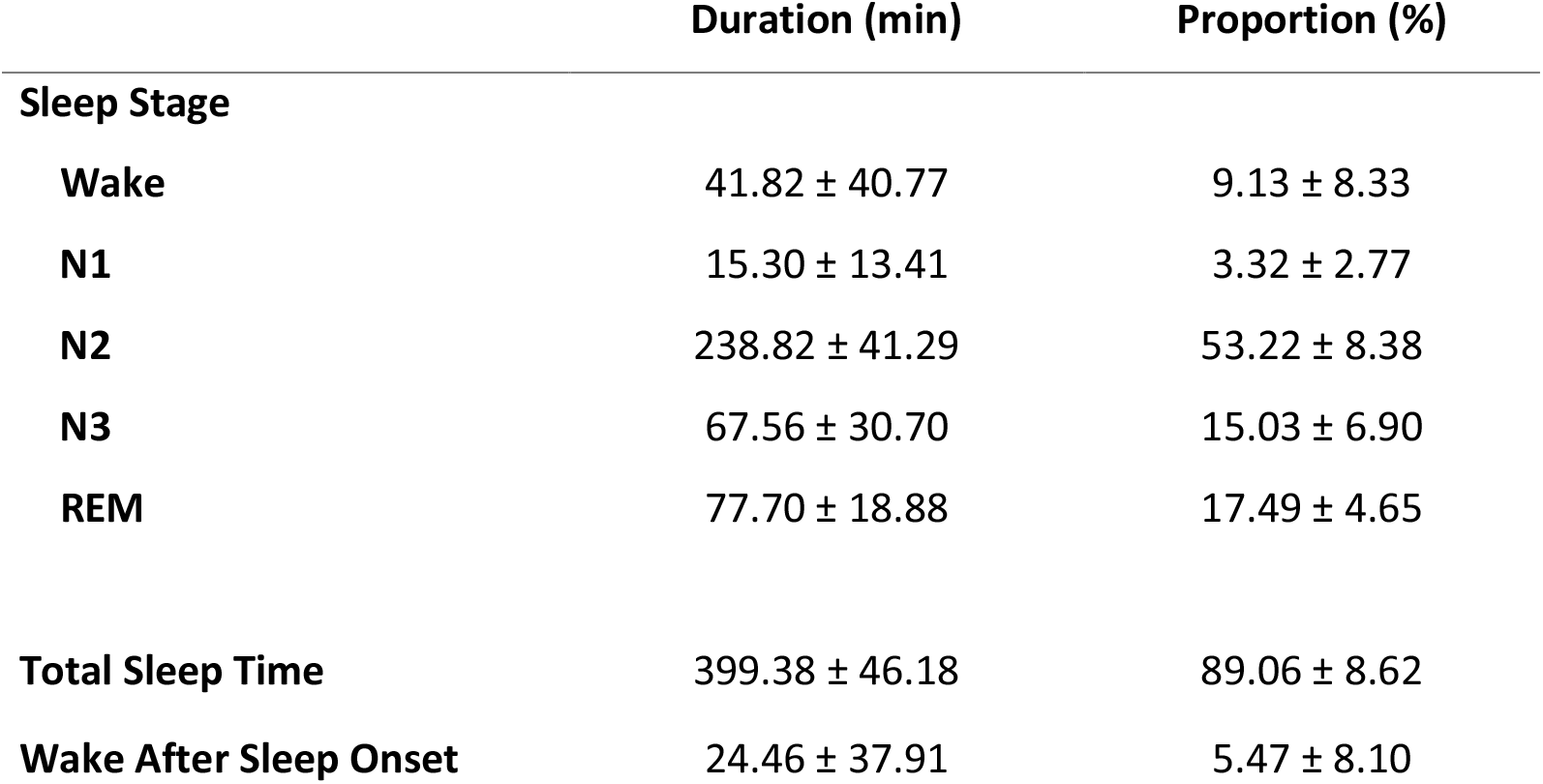
Mean duration (min) and proportion over the night (%) for each sleep stage, Total Sleep Time and Wake After Sleep Onset (mean ± SD).

|  | Duration (min) | Proportion (%) |
| --- | --- | --- |
| **Sleep Stage** |  |  |
| **Wake** | 41.82 ± 40.77 | 9.13 ± 8.33 |
| **N1** | 15.30 ± 13.41 | 3.32 ± 2.77 |
| **N2** | 238.82 ± 41.29 | 53.22 ± 8.38 |
| **N3** | 67.56 ± 30.70 | 15.03 ± 6.90 |
| **REM** | 77.70 ± 18.88 | 17.49 ± 4.65 |
| **Total Sleep Time** | 399.38 ± 46.18 | 89.06 ± 8.62 |
| **Wake After Sleep Onset** | 24.46 ± 37.91 | 5.47 ± 8.10 |

**Supplementary Table 2.** Number (#) and proportion (%) of cues associated to an SO, spindle or SO–spindle complex (mean ± SEM).

|  | **SO** |  | **Spindle** |  | **SO–spindle Complex** |  |
| --- | --- | --- | --- | --- | --- | --- |
|  | # | % | # | % | # | % |
| **All** | 187 ± 10.18 | 39.59 ± 1.78 | 271 ± 10.66 | 57.33 ± 1.16 | 115 ± 8.59 | 24.38 ± 1.48 |
| **In-phase** | 92.3 ± 5.51 | 39.15 ± 1.97 | 135 ± 5.81 | 57.07 ± 1.37 | 56.6 ± 4.60 | 23.88 ± 1.66 |
| **Encoded** | 63.6 ± 4.18 | 40.31 ± 2.20 | 91.2 ± 4.26 | 57.61 ± 1.56 | 39.7 ± 3.66 | 24.98 ± 1.96 |
| **Control** | 28.7 ± 1.59 | 36.80 ± 1.90 | 44.1 ± 1.71 | 55.98 ± 1.27 | 16.9 ± 1.28 | 21.65 ± 1.53 |
| **Antiphase** | 94.2 ± 5.27 | 40.01 ± 1.86 | 136 ± 5.25 | 57.56 ± 1.23 | 58.9 ± 4.36 | 24.86 ± 1.49 |
| **Encoded** | 63.8 ± 3.80 | 40.69 ± 2.02 | 91 ± 3.47 | 57.86 ± 1.19 | 39.8 ± 3.04 | 25.23 ± 1.57 |
| **Control** | 30.4 ± 1.64 | 38.68 ± 1.80 | 45.1 ± 1.93 | 56.97 ± 1.64 | 19.1 ± 1.46 | 24.11 ± 1.58 |

**Supplementary Table 3.** Overview of memory performance. Associative memory % refers to the percentage of correctly recalled images (relative to the total number of stimuli). Statistical differences between conditions (in-phase vs antiphase) were assessed using dependent samples t-tests (two-sided).

|  | In-phase | Antiphase | *t* | *p* |
| --- | --- | --- | --- | --- |
| **Recognition [Hits] %** |  |  |  |  |
| **pre-sleep** | 80.00±3.02 | 81.33±3.12 | -0.31 | 0.76 |
| **post-sleep** | 86.40±2.64 | 83.20±3.62 | 0.71 | 0.48 |
| **post- relative to pre-sleep** | 109.61±3.10 | 102.43±2.62 | 1.77 | 0.08 |
| **Associative Memory %** |  |  |  |  |
| **pre-sleep** | 67.47±4.25 | 67.20±4.17 | 0.04 | 0.96 |
| **post-sleep** | 68.67±4.24 | 64.80±4.50 | 0.63 | 0.53 |
| **post- relative to pre-sleep** | 102.12±1.45 | 95.84±1.74 | 2.77 | **0.01** |

## References

1. Brodt, S., Inostroza, M., Niethard, N. & Born, J. Sleep—A brain-state serving systems memory consolidation. Neuron 111, 1050–1075 (2023).

2. Klinzing, J. G., Niethard, N. & Born, J. Mechanisms of systems memory consolidation during sleep. Nat Neurosci 22, 1598–1610 (2019).

3. Girardeau, G. & Lopes-dos-Santos, V. Brain neural patterns and the memory function of sleep. Science 374, 560–564 (2021).

4. Navarrete, M., Valderrama, M. & Lewis, P. A. The role of slow-wave sleep rhythms in the cortical-hippocampal loop for memory consolidation. Current Opinion in Behavioral Sciences 32, 102–110 (2020).

5. Schreiner, T. & Staudigl, T. Electrophysiological signatures of memory reactivation in humans. Phil. Trans. R. Soc. B 375, 20190293 (2020).

6. Amzica, F. & Steriade, M. The functional significance of K-complexes. Sleep Med Rev 6, 139–149 (2002).

7. Isomura, Y. et al. Integration and segregation of activity in entorhinal-hippocampal subregions by neocortical slow oscillations. Neuron 52, 871–882 (2006).

8. Steriade, M., McCormick, D. A. & Sejnowski, T. J. Thalamocortical oscillations in the sleeping and aroused brain. Science 262, 679–685 (1993).

9. Kim, J., Gulati, T. & Ganguly, K. Competing Roles of Slow-Oscillations and Delta-Waves in Memory Consolidation Versus Forgetting. Cell 179, 514–526.e13 (2019).

10. Muehlroth, B. E. et al. Precise Slow Oscillation–Spindle Coupling Promotes Memory Consolidation in Younger and Older Adults. Sci Rep 9, 1940 (2019).

11. Niethard, N., Ngo, H.-V. V., Ehrlich, I. & Born, J. Cortical circuit activity underlying sleep slow oscillations and spindles. Proceedings of the National Academy of Sciences 115, E9220–E9229 (2018).

12. Maingret, N., Girardeau, G., Todorova, R., Goutierre, M. & Zugaro, M. Hippocampo-cortical coupling mediates memory consolidation during sleep. Nat Neurosci 19, 959–964 (2016).

13. Schreiner, T. et al. Spindle-locked ripples mediate memory reactivation during human NREM sleep. Nat Commun 15, 5249 (2024).

14. Schwimmbeck, F. et al. Sequential coupling of sleep oscillations enables concept-neuron reactivation and supports information flow across the human hippocampal-cortical circuit. Preprint at 10.64898/2026.01.15.699122 (2026).

15. Schreiner, T., Petzka, M., Staudigl, T. & Staresina, B. P. Endogenous memory reactivation during sleep in humans is clocked by slow oscillation-spindle complexes. Nat Commun 12, 3112 (2021).

16. Hahn, M. A., Heib, D., Schabus, M., Hoedlmoser, K. & Helfrich, R. F. Slow oscillation-spindle coupling predicts enhanced memory formation from childhood to adolescence. Elife 9, e53730 (2020).

17. Helfrich, R. F., Mander, B. A., Jagust, W. J., Knight, R. T. & Walker, M. P. Old Brains Come Uncoupled in Sleep: Slow Wave-Spindle Synchrony, Brain Atrophy, and Forgetting. Neuron 97, 221–230.e4 (2018).

18. Mikutta, C. et al. Phase-amplitude coupling of sleep slow oscillatory and spindle activity correlates with overnight memory consolidation. Journal of Sleep Research 28, e12835 (2019).

19. Mölle, M., Yeshenko, O., Marshall, L., Sara, S. J. & Born, J. Hippocampal sharp wave-ripples linked to slow oscillations in rat slow-wave sleep. J Neurophysiol 96, 62–70 (2006).

20. Mölle, M., Eschenko, O., Gais, S., Sara, S. J. & Born, J. The influence of learning on sleep slow oscillations and associated spindles and ripples in humans and rats. Eur J Neurosci 29, 1071–1081 (2009).

21. Staresina, B. P. et al. Hierarchical nesting of slow oscillations, spindles and ripples in the human hippocampus during sleep. Nat Neurosci 18, 1679–1686 (2015).

22. Corzo, A. S. et al. Respiratory coordination of excitability states across the human wake-sleep cycle. Prog Neurobiol 256, 102857 (2026).

23. Heck, D. H., et al. Breathing as a Fundamental Rhythm of Brain Function. Front. Neural Circuits 10, (2017).

24. Kluger, D. S. et al. Modulatory dynamics of periodic and aperiodic activity in respiration-brain coupling. Nat Commun 14, 4699 (2023).

25. Tort, A. B. L., Laplagne, D. A., Draguhn, A. & Gonzalez, J. Global coordination of brain activity by the breathing cycle. Nat. Rev. Neurosci. 1–21 (2025) doi:10.1038/s41583-025-00920-7.

26. Casali, G. et al. Respiratory pauses highlight sleep architecture in mice. Nat Commun 17, 6620 (2026).

27. Brændholt, M. et al. The respiratory cycle modulates distinct dynamics of affective and perceptual decision-making. PLOS Computational Biology 21, e1013086 (2025).

28. Bullón Tarrasó, E., et al. Respiration Shapes the Neural Dynamics of Successful Remembering in Humans. J. Neurosci. 46, e1221252025 (2026).

29. Heck, D. H., Kozma, R. & Kay, L. M. The rhythm of memory: how breathing shapes memory function. Journal of Neurophysiology 122, 563–571 (2019).

30. Zelano, C. et al. Nasal Respiration Entrains Human Limbic Oscillations and Modulates Cognitive Function. J. Neurosci. 36, 12448–12467 (2016).

31. Ghibaudo, V., Juventin, M., Buonviso, N. & Peter-Derex, L. The timing of sleep spindles is modulated by the respiratory cycle in humans. Clin Neurophysiol 166, 252–261 (2024).

32. Schreiner, T., Petzka, M., Staudigl, T. & Staresina, B. P. Respiration modulates sleep oscillations and memory reactivation in humans. Nat Commun 14, 8351 (2023).

33. Schwimmbeck, F., Bullón Tarrasó, E. & Schreiner, T. A role for respiration in coordinating sleep oscillations and memory consolidation. Trends in Neurosciences 0, (2025).

34. Hu, X., Cheng, L. Y., Chiu, M. H. & Paller, K. A. Promoting memory consolidation during sleep: A meta-analysis of targeted memory reactivation. Psychological Bulletin 146, 218–244 (2020).

35. Krugliakova, E. et al. Hacking the functions of sleep: noninvasive approaches to stimulate sleep neurophysiology. Physiological Reviews 106, 675–749 (2026).

36. Ngo, H.-V. V., Martinetz, T., Born, J. & Mölle, M. Auditory Closed-Loop Stimulation of the Sleep Slow Oscillation Enhances Memory. Neuron 78, 545–553 (2013).

37. Sheriff, A. et al. Breathing orchestrates synchronization of sleep oscillations in the human hippocampus. Proceedings of the National Academy of Sciences 121, e2405395121 (2024).

38. Cairney, S. A., Guttesen, A. Á. V., El Marj, N. & Staresina, B. P. Memory Consolidation Is Linked to Spindle-Mediated Information Processing during Sleep. Curr Biol 28, 948–954.e4 (2018).

39. Schechtman, E. et al. Multiple memories can be simultaneously reactivated during sleep as effectively as a single memory. Commun Biol 4, 25 (2021).

40. Kerrén, C., Linde-Domingo, J., Hanslmayr, S. & Wimber, M. An Optimal Oscillatory Phase for Pattern Reactivation during Memory Retrieval. Current Biology 28, 3383–3392.e6 (2018).

41. King, J.-R. & Dehaene, S. Characterizing the dynamics of mental representations: the temporal generalization method. Trends in Cognitive Sciences 18, 203–210 (2014).

42. Özkurt, T. E. & Schnitzler, A. A critical note on the definition of phase-amplitude cross-frequency coupling. J Neurosci Methods 201, 438–443 (2011).

43. Linde-Domingo, J., Treder, M. S., Kerrén, C. & Wimber, M. Evidence that neural information flow is reversed between object perception and object reconstruction from memory. Nat Commun 10, 179 (2019).

44. Farthouat, J., Gilson, M. & Peigneux, P. New evidence for the necessity of a silent plastic period during sleep for a memory benefit of targeted memory reactivation. Sleep Spindles & Cortical Up States 1, 14–26 (2017).

45. Schreiner, T., Lehmann, M. & Rasch, B. Auditory feedback blocks memory benefits of cueing during sleep. Nat Commun 6, 8729 (2015).

46. Karalis, N. & Sirota, A. Breathing coordinates cortico-hippocampal dynamics in mice during offline states. Nat Commun 13, 467 (2022).

47. Harrington, M. O. & Cairney, S. A. Sounding It Out: Auditory Stimulation and Overnight Memory Processing. Curr Sleep Medicine Rep 7, 112–119 (2021).

48. Ngo, H.-V. V. & Staresina, B. P. Shaping overnight consolidation via slow-oscillation closed-loop targeted memory reactivation. Proceedings of the National Academy of Sciences 119, e2123428119 (2022).

49. Bendor, D. & Wilson, M. A. Biasing the content of hippocampal replay during sleep. Nat Neurosci 15, 1439–1444 (2012).

50. Rothschild, G., Eban, E. & Frank, L. M. A cortical-hippocampal-cortical loop of information processing during memory consolidation. Nat Neurosci 20, 251–259 (2017).

51. Schreiner, T., Doeller, C. F., Jensen, O., Rasch, B. & Staudigl, T. Theta Phase-Coordinated Memory Reactivation Reoccurs in a Slow-Oscillatory Rhythm during NREM Sleep. Cell Rep 25, 296–301 (2018).

52. Buzsáki, G. Hippocampal sharp wave-ripple: A cognitive biomarker for episodic memory and planning. Hippocampus 25, 1073–1188 (2015).

53. Sirota, A., Csicsvari, J., Buhl, D. & Buzsáki, G. Communication between neocortex and hippocampus during sleep in rodents. Proc Natl Acad Sci U S A 100, 2065–2069 (2003).

54. Liu, Y., McAfee, S. S. & Heck, D. H. Hippocampal sharp-wave ripples in awake mice are entrained by respiration. Sci Rep 7, 8950 (2017).

55. Buysse, D. J., Reynolds, C. F., Monk, T. H., Berman, S. R. & Kupfer, D. J. The Pittsburgh sleep quality index: A new instrument for psychiatric practice and research. Psychiatry Research 28, 193–213 (1989).

56. Horne, J. A. & Ostberg, O. A self-assessment questionnaire to determine morningness-eveningness in human circadian rhythms. Int J Chronobiol 4, 97–110 (1976).

57. Chung, F., Abdullah, H. R. & Liao, P. STOP-Bang Questionnaire: A Practical Approach to Screen for Obstructive Sleep Apnea. Chest 149, 631–638 (2016).

58. Malach, R. et al. Object-related activity revealed by functional magnetic resonance imaging in human occipital cortex. Proc Natl Acad Sci U S A 92, 8135–8139 (1995).

59. Epstein, R. & Kanwisher, N. A cortical representation of the local visual environment. Nature 392, 598–601 (1998).

60. Konkle, T., Brady, T. F., Alvarez, G. A. & Oliva, A. Conceptual distinctiveness supports detailed visual long-term memory for real-world objects. Journal of Experimental Psychology: General 139, 558–578 (2010).

61. Renard, Y., et al. OpenViBE: An Open-Source Software Platform to Design, Test, and Use Brain–Computer Interfaces in Real and Virtual Environments. Presence: Teleoperators and Virtual Environments 19, 35–53 (2010).

62. Iber, C., Ancoli-Israel, S., Chesson, A. L. & Quan, S. F. The new sleep scoring manual - The evidence behind the rules. Journal of Clinical Sleep Medicine 3, 107 (2007).

63. Grootswagers, T., Wardle, S. G. & Carlson, T. A. Decoding Dynamic Brain Patterns from Evoked Responses: A Tutorial on Multivariate Pattern Analysis Applied to Time Series Neuroimaging Data. J Cogn Neurosci 29, 677–697 (2017).

64. Berens, P. Circular Statistics Toolbox (Directional Statistics). MATLAB Central File Exchange (2026).

65. Oostenveld, R., Fries, P., Maris, E. & Schoffelen, J.-M. FieldTrip: Open Source Software for Advanced Analysis of MEG, EEG, and Invasive Electrophysiological Data. Computational Intelligence and Neuroscience 2011, 1–9 (2011).

